# Deconvoluting signals downstream of growth and immune receptor kinases by phosphocodes of the BSU1 family phosphatases

**DOI:** 10.1101/685610

**Authors:** Chan Ho Park, Yang Bi, Ji-Hyun Youn, So-Hee Kim, Jung-Gun Kim, Nicole Y. Xu, Ruben Shrestha, Alma L Burlingame, Shou-Ling Xu, Mary Beth Mudgett, Seong-Ki Kim, Tae-Wuk Kim, Zhi-Yong Wang

## Abstract

Hundreds of leucine-rich repeat receptor kinases (LRR-RKs) have evolved to control diverse processes of growth, development, and immunity in plants; the mechanisms that link LRR-RKs to distinct cellular responses are not understood. Here we show that two LRR-RKs, the brassinosteroid hormone receptor BRI1 (BRASSINOSTEROID INSENSITIVE 1) and the flagellin receptor FLS2 (FLAGELLIN SENSING 2), regulate downstream glycogen synthase kinase 3 (GSK3) and mitogen-activated protein (MAP) kinases, respectively, through phosphocoding of the BRI1-SUPPRESSOR1 (BSU1) phosphatase. BSU1 was previously identified as a component that inactivates GSK3s in the BRI1 pathway. We found surprisingly that loss of the BSU1 family phosphatases activates effector-triggered immunity (ETI) and impairs flagellin-triggered MAP kinase activation and immunity. The flagellin-activated BOTRYTIS-INDUCED KINASE 1 (BIK1) phosphorylates BSU1 at serine-251. Mutation of serine-251 reduces the ability of BSU1 to mediate flagellin-induced MAP kinase activation and immunity, but not its abilities to suppress ETI and interact with GSK3, which is enhanced through the phosphorylation of BSU1 at serine-764 upon brassinosteroid signaling. These results demonstrate that BSU1 plays an essential role in immunity and transduces brassinosteroid-BRI1 and flagellin-FLS2 signals using different phosphorylation sites. Our study illustrates that phosphocoding in shared downstream components provides signaling specificities for diverse plant receptor kinases.

---

The gene family of leucine-rich repeat receptor kinases (LRR-RK) has expanded dramatically in higher plants, with over 200 and 400 members in *Arabidopsis* and rice, respectively^1^. Among the *Arabidopsis* LRR-RKs, the brassinosteroid (BR) receptor BRI1 and the flagellin receptor FLS2 have distinct functions but are mechanistically related. Activation of BRI1 by BR and activation of FLS2 by the pathogen-associated molecular pattern (PAMP) flagellin involve ligand-induced association with the common co-receptor BRI1-ASSOCIATED KINASE1 (BAK1)^2-4^. Signal transduction downstream of BRI1 is mediated by a well-characterized cascade of phospho-relay events, which include BRI1-mediated phosphorylation of the BR-SIGNALING KINASE (BSK) and CONSTITUTIVE DIFFERENTIAL GROWTH 1 (CDG1) family of receptor-like cytoplasmic kinases (RLCKs)^5^, CDG1-mediated phosphorylation of serine-764 (S764) of the BRI1-SUPPRESSOR1 (BSU1) phosphatase^6^, and BSU1-mediated dephosphorylation of the GSK3 kinase BRASSINOSTEROID-INSENSITIVE 2 (BIN2)^7^, leading to SERINE/THREONINE PROTEIN PHOSPHATASE 2A (PP2A)-mediated dephosphorylation of the BRASSINAZOLE-RESISTANT 1 (BZR1) family of transcription factors^8,9^. Signaling pathways downstream of FLS2 also involve RLCKs such as BIK1 and PBS1-LIKE (PBL) kinases^10,11^, which activate the MAP kinases through mechanisms that are not fully understood^12^. In addition to sharing the co-receptor kinase BAK1, BRI1 and FLS2 have been shown to also share their substrates BSK1 and BIK1 kinases^13,14^. The molecular mechanisms that ensure signaling specificity of these shared downstream components remain unknown.

Plant immune responses are mediated by two systems: pattern-triggered immunity (PTI) and effector-triggered immunity (ETI). PTI is activated by pattern recognition receptors (PRRs) that recognize PAMPs at the cell surface. Pathogens deliver effector proteins into plant cells to attenuate plant PTI. Plants then evolved intracellular receptors to recognize the perturbations of PTI pathways by pathogen effectors^15^. Recent studies showed that PTI and ETI activate overlapping downstream defense outputs and mutually potentiate each other^16-18^. Genetic perturbations of some PTI components, such as FLS2’s co-receptor BAK1 and substrate BIK1, activate ETI and increase resistance to virulent bacterial pathogens^19,20^.

The *Arabidopsis* BSU1 family of phosphatases includes 4 members named BSU1 and BSU1-LIKE 1 (BSL1) to BSL3 (together named BSLs)^21,22^. In addition to mediating BR signaling, The BSU1 family also plays an essential role in stomatal development^22^. BSLs promote stomatal asymmetric cell division by establishing kinase-based signalling asymmetry in the two daughter cells^34^. The partial loss-of-function quadruple mutant (*bsu1;bsl1;amiBSL2/3*, named *bsu-q*) displays pleiotropic defects and the *bsl1;bsl2,bsl3* triple mutant is embryo lethal, which suggest additional roles beyond BR signaling and stomata development^22,23^.

## Results

### BSU1 family plays an essential role in immunity

To fully understand the functions of the BSU1 family phosphatases, we performed RNA-seq analysis of *bsu-q*. The results show not only decreased expression of growth-related genes as expected but also increased expression of genes related to immunity in *bsu-q* (Supplementary Table 1 and 2; Extended Data Fig. 1).

To understand the function of BSLs in immunity, we analyzed how *bsu-q* affects the expression levels of genes that are activated during ETI^16^ and PTI (flagellin peptide flg22, Supplementary Table 3). Consistent with the GO analysis, more ETI-and flg22-induced genes were increased (178) than decreased (56) in *bsu-q*. Compared to wild type, *bsu-q* increased the expression levels (>2x) of a large portion (82 of 823) of the genes activated by both ETI and flg22, and smaller but significant portions of genes activated by only ETI (56 of 1374) or only flg22 (40 of 547) (Fig. 1a). Many flg22-insensitive but ETI-activated genes were dramatically increased in *bsu-q* (Fig. 1b). In contrast, many ETI-independent but flg22-induced genes, including 11 of the 53 genes induced over 5-fold by flg22 in wild-type plants, showed no flg22 response in *bsu-q* (Fig. 1c). The *bsu-q* mutant showed a reduced flg22 responsiveness at the transcriptomic level (Supplementary Tables 3; Extended data Fig. 2a, b). Quantitative PCR analysis confirmed the RNA-seq results for several selected genes that showed elevated expression levels *(FLG22-INDUCED RECEPTOR-LIKE KINASE 1* (*FRK1*), *At2g17740*, and *CYSTEINE-RICH RLK 13* (*CRK13*)) or insensitivity to flg22 (*ROF2*) in *bsu-q* (Extended data Fig. 2c). These results suggest that the *bsu-q* mutant has a constitutive activation of ETI response and a defect in flagellin signaling. Similar phenotypes have been observed in mutants lacking components of PAMP-signaling, such as the *bak1 bkk1* mutant which lacks the co-receptor for FLS2 and BRI1. Indeed, a comparison of transcriptomes affected in *bak1 bkk1*^19^ with those affected in *bsu-q* revealed that the mutants share similar gene expression patterns (Fig. 1d). Of the 509 genes upregulated in *bsu-q*, 240 (47%) were also upregulated in *bak1 bkk1*, and these include 132 (55%) ETI-and/or flg22-induced genes (Fig. 1d). The transcriptomic analyses support a possibility that BSU1 family phosphatases play dual roles in immunity and BR signaling, similar to that observed for the upstream signaling components BAK1, BIK1, and BSK1^13,14,19^.

**Fig. 1.**
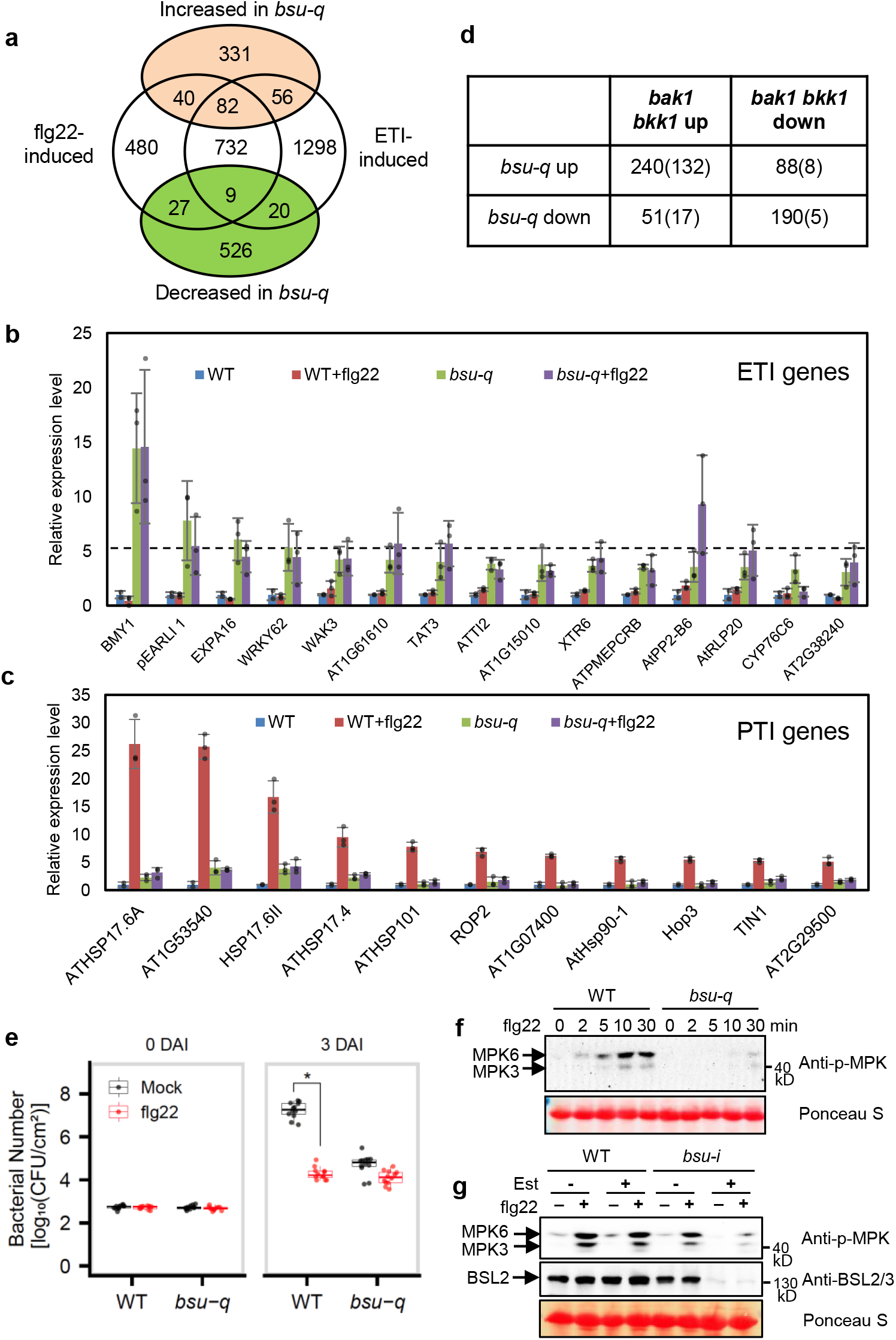
The *bsu-q* mutant shows enhanced ETI and defective flagellin signaling. (a) The Venn diagram shows the overlaps between the genes up-or down-regulated in *bsu-q* relative to wild type and the genes activated by flg22 treatment or ETI. (b-c) Relative expression 478 levels normalized to wild type (WT) of (b) ETI genes that did not respond to flg22 in wild type 479 but increased in *bsu-q*, and (c) 11 genes, among the 53 ETI-independent genes that were induced 480 over 5-fold by flg22 in WT but showed no significant response to flg22 (FC<2) in *bsu-q*. The graph shows mean +/- SEM values from the RNA-seq data in Supplementary Table 4. (d) Overlap between differentially expressed genes of *bsu-q* and *bak1 bkk1*. Numbers in parenthesis are ETI/flg22-responsive genes. (e) Quantitation of *Pseudomonas syringae* pathovar *tomato* strain DC3000 in WT and *bsu-q* plants treated with mock or flg22. Each replicate includes four leaf discs and 12 biological repeats (n=12) were used, and the experiment was repeated three times with similar results. The box plot shows the minimum, first quantile, median, third quantile and maximum of the 12 displayed data points for each sample. Asterisk indicates statistically significant difference by two-tail t-test (p=7.115e-16). (f) The *bsu-q* mutant is defective in flg22-induced MPK3/6 phosphorylation. Leaf tissues of wild type (WT) and *bsu*-*q* were treated with 100 nM flg22 for the indicated time, and MAP kinase activation was analyzed by immunoblot using anti-phospho-p44/42 MAPK antibody. Ponceau S staining of membrane shows equal protein loading. The experiment was repeated three times with similar results.(g) WT and *bsu-i* seedlings were grown on medium containing 0 or 20 μM estradiol for nine days, and then treated with 100 nM flg22 for 10 min. Immunoblots were probed with anti-phospho-p44/42 MAPK and anti-BSL2/3 antibodies. Ponceau S staining of membrane shows equal protein loading. The experiment was repeated twice with similar results.

To test whether the altered defense gene expression affects immune response in *bsu-q*, we analyzed bacterial growth after inoculation of wild type and *bsu-q* mutant with *Pseudomonas syringae* pathovar *tomato* (*Pst*) strain DC3000. The results show that *bsu-q* is more resistant than wild type to *Pst* infection (Fig. 1e). Pretreatment of seedlings with flg22 inhibited bacterial growth in the wild-type plants but did not further enhance disease resistance in the *bsu-q* mutant (Fig. 1e).

The priming of defense responses in untreated *bsu-q* plants and compromised flg22-triggered responses are consistent with the altered expression of ETI and PTI genes. These phenotypes are similar to those of other mutants with loss of PAMP signaling components, including the *bik1*^20^ and *bak1 bkk1* double mutant^19^. The *bak1 bkk1* double mutant exhibits impaired PTI responses such as flagellin-induced MAP kinase activation and reactive oxygen response (ROS), as well as enhanced ETI and accumulation of the defense hormone salicylic acid (SA)^19^. Similarly, we observed a weaker flg22-induced ROS response in the *bsu-q* mutant and an elevated level of SA in the inducible quadruple mutant *bsu-i* plants (*bsu1;bsl1;bsl3* triple mutant transformed with a construct for estradiol-inducible expression of an artificial microRNA targeting *BSL2*. See Methods) (Extended Data Fig. 3). These defense-related phenotypes similar to the *bak1 bkk1* and *bik1* mutants further support the hypothesis that BSU1 family members have dual functions, similar to BAK1 and BIK1, as components of PTI signaling pathways and guardees of resistance proteins operating in ETI signaling pathways. In line with this hypothesis, a recent study identified the BSU1 family members as targets of pathogen effector proteins^24^.

### BSU1 mediates flagellin signaling as a BIK1 kinase substrate

To determine whether the BSU1 family members play a role in FLS2 signaling, we analyzed flg22-induced activation of MAP kinases in mutants lacking various BSU1/BSLs. Flg22 treatment induced similar MAP kinase phosphorylation in wild type and *bsl2;bsl3* double mutant (Extended Data Fig. 4a), but induced much weaker MAP kinase phosphorylation in the *bsu-q* mutant (Fig. 1f) and in the *bsu-i* plants (Fig. 1g), indicating that BSU1/BSLs are redundantly required for flg22-induced MAP kinase activation. In addition, MAP kinase phosphorylation caused by another bacterial elicitor, elf18^25^, was reduced in the estradiol-treated *bsu-i* mutant (Extended Data Fig. 4b), suggesting a more general role for BSU1/BSLs in PTI signaling.

The mode of function of BSU1 in the BR signaling pathway suggests that BSU1 may transduce the flagellin signal from BIK1 to downstream MAP kinases. We thus performed yeast-two-hybrid assays to test if BSU1/BSLs interact with BIK1 and MEKK1, a MAP kinase kinase kinase (MAPKKK) involved in flagellin-induced MAP kinase activation^26,27^. The results show that BSU1 interacts with both BIK1 and MEKK1 (Fig. 2a,b) and that BSU1/BSLs interact with multiple BIK1/PBL proteins with various specificities and affinities (Extended Data Fig. 5), suggesting that different BSU1 and BIK1/PBL family members may act redundantly in parallel pathways. Consistent with the yeast two-hybrid assay results, *in vitro* pull-down assays showed that BIK1 interacts with BSU1 but not the BSLs (Fig. 2c), and BSU1 also interacts with MEKK1 *in vitro* (Fig. 2d). Moreover, co-immunoprecipitation assays confirmed the interaction between BIK1-Myc and BSU1-YFP in *Nicotiana benthamiana* and transgenic *Arabidopsis* plants (Fig. 2e), as well as between BSU1-Myc and MEKK1-YFP (Fig. 2f). These results suggest a potential role for BSU1 in transducing the signal from BIK1 to MEKK1.

**Fig. 2.**
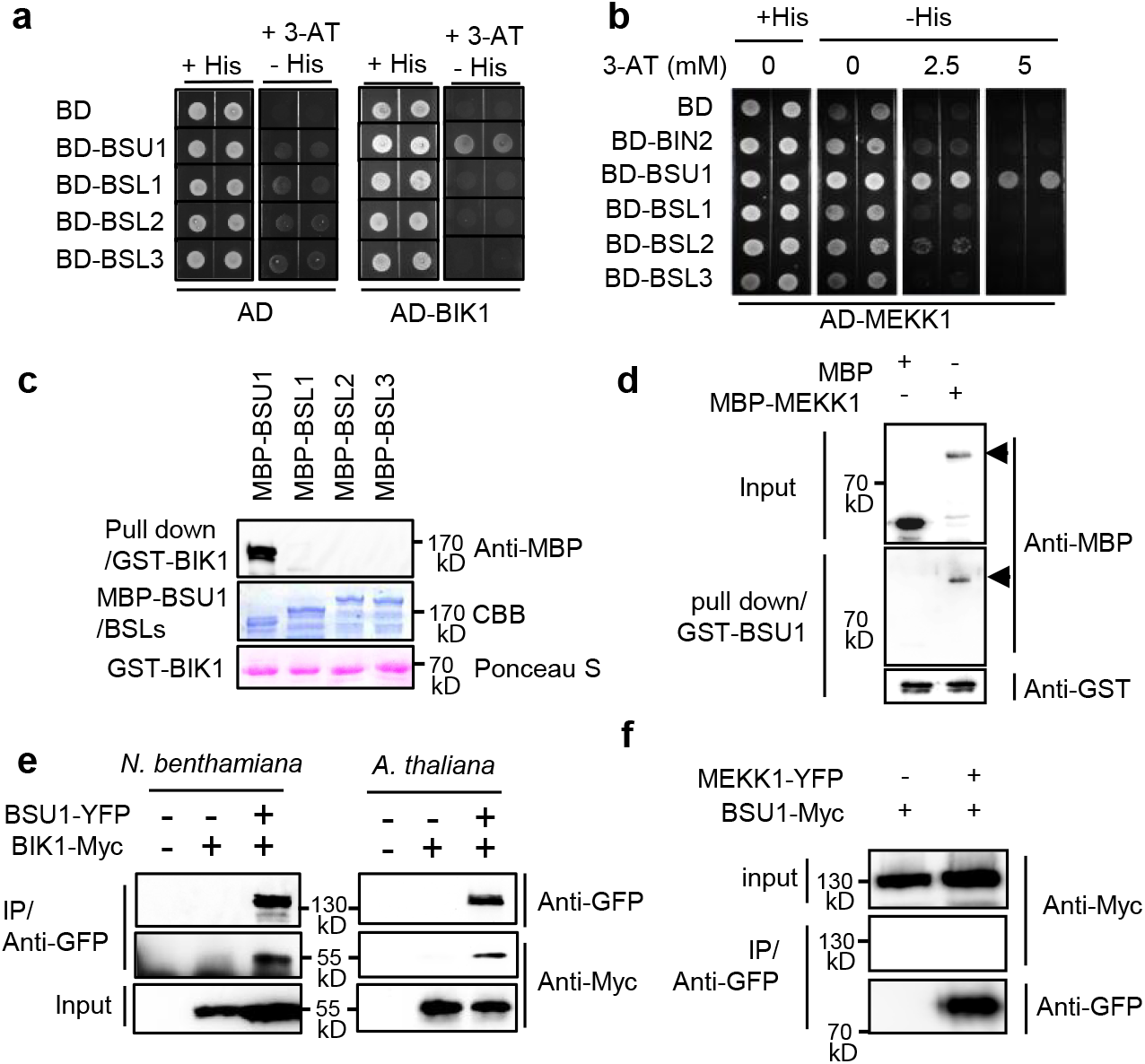
BSU1 interacts with BIK1 and MEKK1. (a) Yeast-two-hybrid assays of interaction between BIK1 and BSU1/BSLs. Yeast cells co-expressing AD-BIK1 and BD-BSU1/BSLs were grown on synthetic dropout medium containing 2.5 mM 3-amino-1, 2, 4-triazole (3-AT). (b) Yeast-two-hybrid assays of interaction between MEKK1 and BSU1/BSLs. (c) *In vitro* pull-down of BSU1/BSLs (as MBP fusions) by GST-BIK1, analyzed by anti-MBP immunoblotting. Input proteins are shown by Coomassie brilliant blue (CBB) of the SDS-PAGE gels and Ponceau S staining of membranes. The experiment was repeated twice with similar results. (d) *In vitro* pull-down of MBP-MEKK1 by GST-BSU1, analyzed by immunoblotting using anti-MBP and anti-GST antibodies. Arrow heads indicate MBP-MEKK1. The experiment was repeated twice with similar results. (e) Co-immunoprecipitation assay of BSU1-YFP with BIK1-Myc protein expressed in *N. benthamiana* leaves and *Arabidopsis* transgenic plants. The experiment was repeated twice in *Arabidopsis* with similar results. (f) Co-immunoprecipitation assay of BSU1-Myc with MEKK1-YFP protein expressed in *N. benthamiana* leaves. The experiment was repeated twice with similar results.

The interaction with BIK1 suggests that BSU1 might be phosphorylated upon flagellin signaling. To test this possibility, we performed metabolic <u>S</u>table Isotope <u>L</u>abeling In *<u>A</u>rabidopsis* followed by Immuno-<u>P</u>recipitation and <u>M</u>ass <u>S</u>pectrometry (SILIA IP-MS) analysis of BSU1-YFP in transgenic *Arabidopsis*. We found that flg22 induced phosphorylation of the serine-251 (S251) residue of BSU1 (Fig. 3a,b). Targeted quantification using parallel reaction monitoring (PRM) mass spectrometry further showed that phosphorylation of BSU1 at S251 was increased over 4 fold after flg22 treatment in both replicate experiments (Fig. 3c,d and Extended Data Fig. 6a,b). The S251 residue of BSU1 is located within the N-terminal Kelch-repeat domain and is conserved in BSL homologs, and paralogs in other plant species (Fig. 3e, Extended Data Fig. 7).

**Fig. 3.**
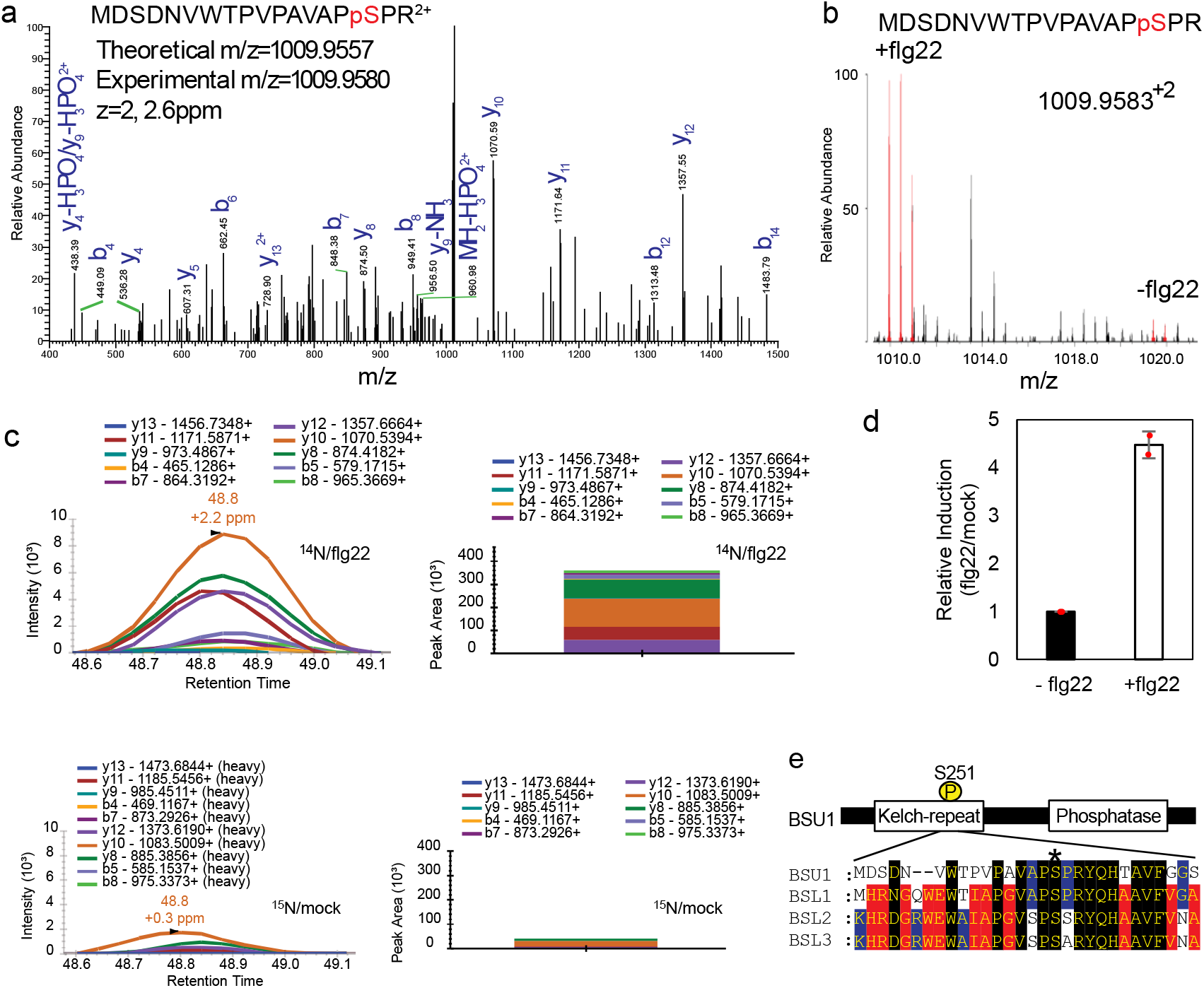
Flagellin induces phosphorylation of BSU1 at S251. (a) MS/MS spectrum of the peptide containing phosphorylated S251 of BSU1. (b) MS1 spectrum of a SILIA IP-MS experiment shows flg22-induced phosphorylation of BSU1 S251. The peaks of ^14^N-labeled (+flg22) and ^15^N-labeled (-flg22) phospho-S251-containing peptides are shown in red. (c) PRM quantitation of the phospho-S251 peptide after SILIA and immunoprecipitation of BSU1-YFP from mock-treated ^15^N-labeled and flg22-treated ^14^N-labeled seedlings. Graphs display chromatograms of fragment ions extracted from the ^14^N-and ^15^N-labeled phospho-peptides. Peak areas of fragment ions are extracted using <5ppm mass window and integrated across the elution profile. (d) Quantification of relative induction (flg22/mock). The graph shows mean ± SEM from two reciprocal SILIA IP-MS experiments (n=2). (e) Alignment of sequences of BSU1 family members around the BSU1 S251 residue (asterisk).

We next tested whether BIK1 phosphorylates BSU1 using *in vitro* kinase assays. As shown in Figure 4a, GST-BIK1 phosphorylated MBP-BSU1 specifically but not the BSLs, consistent with the yeast interaction results (Fig. 2a). GST-BIK1 phosphorylated the N-terminal Kelch domain more strongly than the C-terminal phosphatase domain (Fig. 4b). The MBP-BSU1 protein containing mutation of S251 to alanine (S251A) was less phosphorylated than wild-type BSU1 by GST-BIK1 (Fig. 4c). In contrast, the alanine substitution of the CDG1 phosphorylation site (S764A) had no obvious effect on BIK1 phosphorylation of BSU1 (Fig. 4c). As reported previously^28^, flg22 treatment induced phosphorylation of BIK1, causing a slower mobility in SDS-PAGE gel and enhanced autophosphorylation (Fig. 4d). BIK1-HA immunoprecipitated from the flg22-treated plants showed stronger phosphorylation of and interaction with MBP-BSU1 than that from untreated plants (Fig. 4d,e). The phosphorylation of BSU1 by BIK1-HA was also decreased by the S251A mutation (Fig. 4f). These results demonstrate that BIK1 phosphorylates the S251 residue of BSU1 in response to flagellin-FLS2 signaling.

**Fig. 4.**
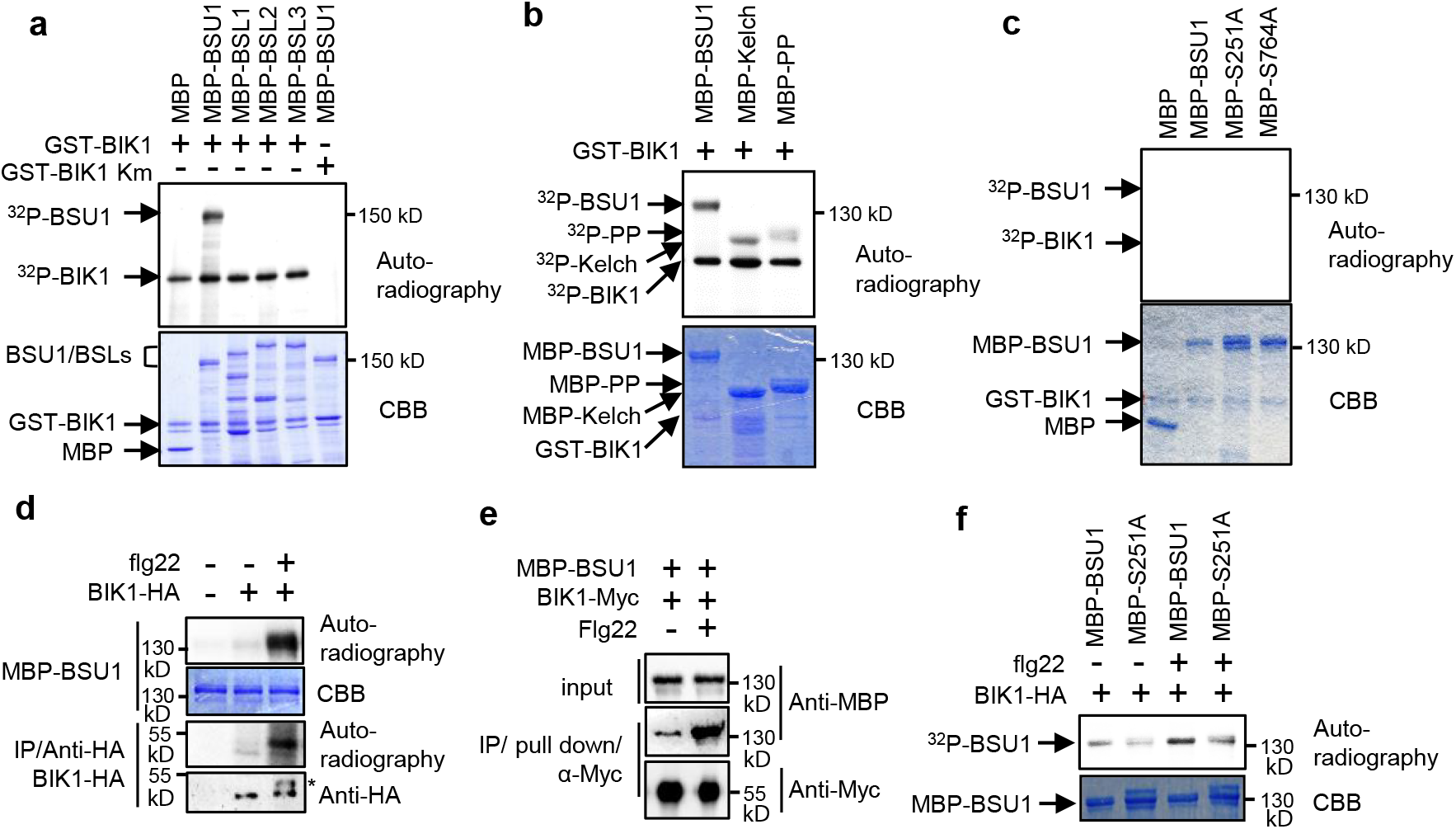
BIK1 phosphorylates BSU1 at S251. (a) *In vitro* kinase assay of BIK1 phosphorylation of the indicated MBP and MBP fusion proteins. Phosphorylation and proteins are shown by autoradiography and gel staining (CBB), respectively. A kinase inactive mutant (Km) BIK1 was used as a negative control. The experiment was repeated three times with similar results. (b) *In vitro* kinase assay of BIK1 phosphorylation of BSU1 N-terminal (Kelch) and C-terminal (PP) domains. The experiment was repeated twice times with similar results. (c) *In vitro* kinase assay of GST-BIK1 phosphorylation of wild-type and mutant (S251A and S764A) MBP-BSU1. The experiment was repeated three times with similar results. (d) *In vitro* kinase assay of flg22-induced BIK1 phosphorylation of BSU1. BIK1-HA was immunoprecipitated using anti-HA antibody from thirteen-day old *BIK1::BIK1-HA* seedlings treated with 1 μM flg22 or mock for 10 min and then incubated with MBP-BSU1 and ^32^P-*γ*-ATP. BIK1-HA proteins were analyzed by immunoblotting using anti-HA antibody. Asterisk indicates phosphorylated BIK1. The experiment was repeated three times with similar results. (e) Flg22-induced BIK1-BSU1 interaction. BIK1-Myc proteins were immuno-purified by anti-Myc antibody from plants treated with 100 nM flg22 or mock for 30 min and were used to pulldown MBP-BSU1 *in vitro*. The experiment was repeated twice with similar results. (f) *In vitro* kinase assay of flg22-induced BIK1 phosphorylation of wild-type and mutant (S251A) BSU1 using immuno-purified BIK1-HA. BIK1-HA was immunoprecipitated using anti-HA antibody from twelve-day old *BIK1::BIK1-HA* seedlings treated with 1 μM flg22 or mock for 10 min, and then incubated with MBP-BSU1 or MBP-BSU1-S251A (MBP-S251A) and ^32^P-*γ*-ATP. The experiment was repeated twice with similar results.

### Phosphorylation of BSU1-S215 mediates MAP kinase activation

To determine whether phosphorylation of S251 in BSU1 plays a role in flagellin-induced MAP kinase activation, we generated transgenic *bsu-i* plants expressing BSU1-YFP or BSU1-S251A-YFP. Two independent transgenic lines expressing wild-type BSU1 in *bsu-i* increased MAP kinase activation by more than 1.7-fold, whereas expression of BSU1-S251A only increased MAP kinase by about 1.3-fold (Fig 5a; Extended Data Fig. 8a).

**Fig. 5.**
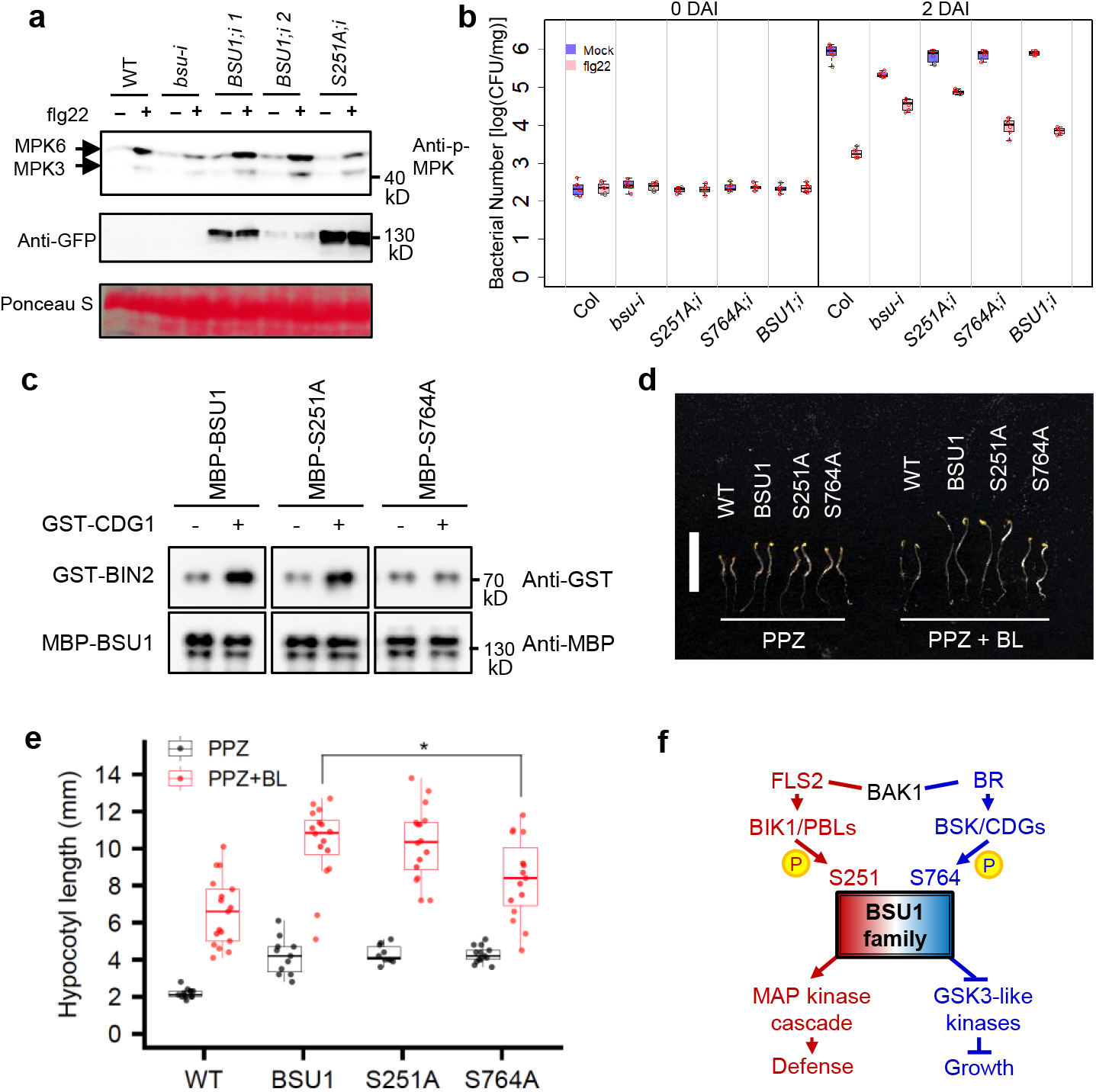
Phosphorylation of BSU1 S251 is required for MAPK activation and immunity but not for BR signaling. (a) Flg22-induced MPK3/6 phosphorylation in *35S::BSU1-YFP;bsu-i* (*BSU1;i*) and 3*5S::BSU1-S251A-YFP;bsu-i* (*S251A;i*) plants. Eight-day old seedlings grown on 20 μM estradiol were treated with 10 nM flg22 for 15 min. Immunoblot was performed using anti-phospho-p44/42 MAPK and anti-GFP antibodies. Ponceau S staining image shows equal loading. The experiment was repeated three times with similar results. (b) Flg22-induced bacterial growth inhibition in seedlings of the indicated genotypes at 0 and 2 days after inoculation (DAI). Each replicate includes 20 mg or more seedlings and 6 biological replicates. The box plot shows the minimum, first quantile, median, third quantile and maximum of the 6 displayed data points for each sample. (c) CDG1-induced BIN2 interaction with MBP-BSU1, MBP-BSU1-S251A and MBP-BSU1-S764A. Wild type and mutant MBP-BSU1 proteins pre-incubated with GST(-) or GST-CDG1(+) were used to pull down GST-BIN2. The experiment was repeated three times with similar results. (d, e) Phenotype (d) and hypocotyl length (e) of 5-day-old etiolated WT, *35S::BSU1-YFP* (BSU1), 3*5S::BSU1-S251A-YFP* (S251A) and 3*5S::BSU1-S764A-YFP* (S764A) seedlings grown on media containing PPZ or PPZ plus BL. The box plot shows the minimum, first quantile, median, third quantile and maximum of the displayed data points for each sample. (d) Bar = 10 mm. (e) The average hypocotyl length ± SEM is shown. Asterisk denotes significant difference from *35S::BSU1-YFP* by two-tailed t-test (p-value =0.01512, n = 16 for BSU1 and n = 15 for S764A). Experiments were performed 3 times and similar results were obtained. (f) Schematic diagram of the BSU1 phosphocodes in BRI1 and FLS2 signaling pathways.

We then tested whether the S251A mutation affects BSU1’s ability to mediate flagellin-induced immunity against pathogen infection. We treated seedlings with mock or flg22 peptide, inoculated them with *Pst* DC3000, and then quantified the number of bacteria in the plant tissue at zero and two days after the inoculation. Flg22 treatment decreased the growth of bacteria by about 464-fold in wild type plants. The mock-treated *bsu-i* plants contained a reduced number of bacteria compared to mock-treated wild type, whereas flg22 treatment caused only about 6-fold reduction of the bacteria growth in *bsu-i*, which was similar to *bsu-q*. These phenotypes are consistent with activation of ETI signaling and a compromise in PTI signaling due to the loss of BSU/BSL family members. The *bsu-i* plants expressing either wild-type BSU1, BSU1-S251A, or BSU1-S764A showed similar pathogen growth as wild type after mock treatment, suggesting that both the wild type and mutant BSU1 proteins rescued the ETI phenotype. Upon flg22 treatment, the *BSU1-251A/bsu-i* plants showed only about 9-fold reduction of pathogen growth, whereas the *BSU1/bsu-i* and *BSU1-S764A/bsu-i* plants showed 110-fold and 72-fold pathogen reduction compared to mock treatment (Fig. 5b). These data indicate that both wild type and mutant BSU1 proteins can suppress the ETI response, whereas BSU1’s function in PTI response is impaired by the S251A mutation but not the S764A mutation. These results further support the conclusions that the presence of BSU1/BSL proteins is required to control ETI, and the BIK1 phosphorylation of BSU1 at S251 contributes to flg22 induction of MAP kinase activation and immunity, whereas phosphorylation of S764 does not play a major role in immune signaling.

Previous studies have shown that BR-BRI1 signaling involves CDG1-mediated phosphorylation of BSU1 S764, which enhances interaction with BIN2, a GSK3-like kinase^6^. To test whether phosphorylation at S251 and S764 specifies downstream targets of BSU1 signaling, we compared the effects of mutations of these residues on BSU1 interaction with BIN2. Consistent with the previous report, phosphorylation by CDG1 increased the interaction of wild-type BSU1 with BIN2 but not that of the BSU1-S764A mutant protein (Fig. 5c). The BSU1-S251A mutant, similar to wild-type BSU1, showed CDG1-induced interaction with BIN2 (Fig. 5c, Extended Data Fig. 8b). Consistent with the normal interaction with BIN2, seedlings expressing BSU1-S251A showed similar BR sensitivity in hypocotyl elongation compared to plants expressing wild type BSU1, whereas plants expressing BSU1-S764A showed reduced sensitivity to BR (Fig. 5d,e, Extended Data Fig. 9). These results show that BSU1’s functions in the flagellin and BR pathways require phosphorylation of distinct residues, S251 and S764, respectively.

### Discussion

Taken together our results show that BSU1 and BSLs redundantly play key roles in plant immunity, in addition to their known functions in BR signaling and stomata development. The presence of a BSU1/BSL protein is required to suppress immune responses and the loss of all BSU1 family members triggers an ETI-like response, whereas flagellin-FLS2 signaling induces BIK1 phosphorylation of BSU1 at S251 which leads to activation of the MAP kinases and immune responses. The mechanism by which phospho-S251 leads to activation of the MAP kinases remains to be investigated. Previous studies have shown that several MEKKs are inhibited by phosphorylation of their N-terminal auto-regulatory domain, suggesting that activation of MEKKs involves dephosphorylation^23,29,30^. Our observations of direct interaction between BSU1 and MEKK1 support a model that BSU1 activates MEKK1 by dephosphorylating its auto-regulatory domain.

BR-BRI1 signaling induces CDG1/BSKs-mediated phosphorylation of BSU1 at S764, which mediates interaction with and dephosphorylation of GSK3-like kinases, leading to growth responses^14^. As such, the signaling specificities for BSU1 in the BRI1 and FLS2 pathways are determined by phosphorylation of different residues, namely phosphocodes (Fig. 5f). Considering the interactions between additional members of the BSU1 and BIK1/PBL families (Extended Data Fig. 5) and the conservation of S251 among BSL family members in *Arabidopsis* and other plants (Extended Data Fig. 7), the mechanism demonstrated here for *Arabidopsis* BSU1 is likely used by its paralogs and homologs in parallel/redundant immune pathways. We propose that phosphocoding, namely distinct signal transduction through specific phosphorylation sites, contributes to the diversification of signaling specificity of the large numbers of receptor kinases in plants. In light of the recent finding of spatial recruitment of BSLs in asymmetric cell division^31^, we propose that phosphocoding and spatial recruitment together provide the signaling specificity that allows the BSU1 family of phosphatases to act in many receptor kinase pathways controlling diverse processes of growth, development, and immunity.

## METHODS

### Plant material and growth condition

All *Arabidopsis thaliana* plants used in this study are in Columbia ecotype background. The *bsu1-1* (SALK_030721)^7^, *bsl1-1* (SALK_051383)^7^, *bsl2-3* (SALK_055335)^22^, *bsl3-2* (SALK_072437)^22^, *bsu-q*^7^, *35S::BSU1-YFP*^7^, *35S::BSU1-Myc*^7^, *BSU1pro::BSU1-YFP*^32^, *BIK1::BIK1-HA*^11^, and *MEKK1-FLAG*^27^ lines were previously described. The *bsl2;bsl3* and *bsu1;bsl1;bsl3* mutants were generated by crossing the single mutants. The estradiol-inducible BSU1 family quadruple mutant (*bsu-i*) was generated by expressing the artificial microRNA targeting BSL2/3 (same as used in *bsu-q*)3 from the estradiol inducible promoter of pMDC7 vector^33^ in the *bsu1;bsl1;bsl3* triple mutant background. *Arabidopsis* seedlings were grown on 0.8% phytoblend agar medium containing 1/2 Murashige-Skoog (MS) basal nutrient and 1% sucrose under continuous light in a growth chamber (Percival) at 22 °C. For estradiol treatment, seeds were planted on the MS medium containing β-estradiol (Sigma E2758). For the RNA-seq experiment, seeds were germinated in the dark for 3 days, seedlings with weak phenotypes (long hypocotyl) were removed from the *bsu-q* samples, and then the plants with strong dwarf phenotypes were grown under light for 12 days. For hypocotyl elongation assay, seeds were planted on MS media containing 2 μM PPZ (Propiconazole, a BR biosynthetic inhibitor, Martin’s HONOR GUARD PPZ)^34^ or 2 μM PPZ and 1 nM BL (Cayman, 21594), and then stratified for 2 days. After stratification, the seeds were treated with light for 6 hr and then incubated in the dark for 5 days. For general growth and seed harvesting, plants were grown in soil in a greenhouse with a 16-hr light/8-hr dark cycle at 22–24 °C.

### Plasmid constructions

To generate the yeast-two-hybrid, *35S::BIK1-Myc, 35S::MEKK1-YFP* and *MBP-MEKK1* constructs, the full-length coding sequences (CDS) without stop codon of the genes were amplified by PCR from *Arabidopsis* cDNA using gene-specific primers described in Supplementary Table 6. Each CDS was cloned into pENTR/SD/D-TOPO vector (Invitrogen) and sub-cloned into Gateway-compatible destination vectors (pCAMBIA1390-4Myc-6His, pEarleyGate101^35^, gcpMALc2x, pXDGATcy86, and pGADT7) by LR reaction. The plasmids for MBP-BSU1, MBP-BSU1-S764A, MBP-BSU1-Kelch, MBP-BSU1-PP, MBP-BSL1, MBP-BSL2, MBP-BSL3, GST-CDG1, GST-BIK1, GST-BIK1Km, 35S::BSU1-YFP, and 35S::BSU1-S764A-YFP were previously described^6,7,10^. The BSU1 clone in pENTR/SD/D-TOPO vector^6^ was used as a template for site-directed mutagenesis following the manufacturer’s protocol (New England Biolab) to generate *BSU1-S251A*, which was subcloned into the Gateway-compatible pEarleyGate101 vector to generate the *35S::BSU1-S251A-YFP* construct.

### Transgenic plants

The *35S::BSU1-S251A-YFP* and *35S::BSU1-S764A-YFP* plasmids were transformed into wildtype *Arabidopsis* using the *Agrobacterium*-mediated floral dipping transformation method. Transgenic lines expressing similar levels of the protein were selected for experiments shown in Fig. 5d-e and Extended Data Fig. 9. The estradiol-inducible BSU1 family quadruple mutant (*bsu-i*) was generated by expressing the artificial microRNA targeting BSL2 (same as used in *bsu-q*)^3^ from the estradiol inducible promoter of the pMDC7 vector^38^ in the *bsu1;bsl1;bsl3* triple mutant. The *35S::BSU1-YFP;bsu-i* (*BSU1;i*), 3*5S::BSU1-S251A-YFP;bsu-i* (*S251A;i*), and 3*5S::BSU1-S764A-YFP;bsu-i* (*S764A;i*) plants were generated by transforming the *bsu-i* plants with the corresponding plasmids.

### PAMP treatment

The flg22 and elf18 peptides were synthesized by the Protein and Nucleic Acid Facility at Stanford University (http://pan.stanford.edu/). The flg22 or elf18 peptides were dissolved in autoclaved distilled water containing 0.05% Silwet L-77 and applied to seedlings or rosette leaves. For RNA-seq experiments, seedlings on plates were submerged in 1 μM flg22 or mock solution for 1 min. After decanting the solution, the seedlings were then incubated in a growth chamber for 1 hr. For Fig. 1f, rosette leaves of eight-week-old short-day-grown plants were treated with 100 nM flg22. For Fig. 1g, nine-day-old seedlings grown on MS media containing 20 μM estradiol were treated with 100 nM flg22 for 10 min. For Fig. 5a, eight-day-old seedlings grown on MS media containing 20 μM estradiol were treated with 10 nM flg22 for 15 min. For Fig. 4d, f, the *BIK1::BIK1-HA* seedlings were treated with 1 μM flg22 for 10 min.

### RNA-seq analysis

Total RNA was extracted from the wild-type and *bsu-q* seedlings treated with flg22 or mock solution using the Spectrum Plant Total RNA kit (Sigma). cDNA libraries were constructed using a TruSeq RNA sample preparation kit (Illumina) according to the manufacturer’s instructions. Samples of three replicates were sequenced on an Illumina HiSEQ2000 sequencing machine. Mapping of reads to the reference genome (TAIR10) and calling differentially expressed genes was conducted via the iPlant collaborative Discovery Environment (de.iplantcollaborative.org/de/)^36^. Expression data was clustered and visualized with the TIGR MeV program (http://mev.tm4.org/)^37^. Scatter plots of flg22-responsive genes were analyzed with Microsoft Excel. GO analysis was performed with agriGO v2.0 (www.http://systemsbiology.cau.edu.cn/agriGOv2/index.php) using default setting^38^.

### Immunoblot analysis

Total proteins from plant materials were extracted with IP buffer (50 mM Tris at pH 7.5, 50 mM NaCl, 0.3 M sucrose, 1% Triton X-100, 1 mM phenylmethylsulfonyl fluoride and 1× protease inhibitor cocktail) or 2× SDS sample buffer (100 mM Tris at pH 6.8, 4% SDS, 20% glycerol and 4% β-mercaptoethanol). After boiling with 2× SDS sample buffer, proteins were loaded on SDS-PAGE gel. Separated proteins were transferred to a nitrocellulose membrane, and probed with anti-phospho-p44/42 mitogen-activated protein kinase (Erk1/2) (Thr202/Tyr204) (Cell Signaling; 1:1,000 dilution), anti-MBP (New England Biolab; 1:2,000 dilution), anti-Myc (Cell Signaling; 1:3,000 dilution), anti-HA (Roche; 1:1,000 dilution), anti-BSL2/3 (AbFrontier; 1:2,000 dilution), anti-FLAG (Sigma; 1:1,000 dilution), anti-GST (Santa Cruz Biotechnology; 1:1,000 dilution) and anti-GFP (homemade or TRANSGENE, M018) antibodies.

### Bacterial growth assays

Bacterial growth assay was performed following published procedures^39^. To monitor *Pseudomonas syringae* pathovar *tomato* (*Pst*) strain DC3000 growth in plants, leaves of 6-week-old plants were hand-infiltrated with a 1×10^5^ colony forming unit (CFU)/mL suspension of the bacteria in 1 mM MgCl2 using a needleless syringe. For flg22 pretreatment, the leaves were hand-infiltrated with water or 1 μM flg22 one day before the bacterial inoculation. Leaf discs were collected and ground in 1 mM MgCl2 and then spotted on nutrient yeast glycerol agar plates in triplicate to determine the bacterial titer. Seedling flood inoculation assay was performed following published procedures with minor modifications^40^. Seedlings grown on media containing 10 μM estradiol were treated with 1 μM flg22 or distilled water one day prior to *Pseudomonas* inoculation. Bacteria inoculum (10^5^ CFU/mL) containing 0.025% Silwet L-77 was dispensed into 7-day-old *Arabidopsis* seedlings grown on plates. Bacterial populations were evaluated from six biological replicates and each replicate represented a pooled sample of at least 20 mg of seedlings.

### Yeast two-hybrid assays and *in vitro* pull-down assay

Yeast two-hybrid assays were performed following published procedures^7^. Recombinant GST-tagged proteins were purified from *E. coli* using Glutathione Sepharose beads (GE healthcare). The GST-tagged protein-bead complexes were incubated with 2 μg of MBP-tagged proteins in 1 mL of pull-down buffer (50 mM Tris, pH 7.5, 150 mM NaCl, 0.2% NP-40 and 0.05 mg/mL BSA) at 4°C for 1 hr. Then, the protein-bead complex was washed twice with 1 mL of pull-down washing buffer (50 mM Tris, pH 7.5, 100 mM NaCl, 0.1% NP-40). Associated proteins were eluted with 2× SDS sample buffer and analyzed by immunoblotting.

### *In vitro* kinase assays

*In vitro* kinase assays using recombinant proteins were performed following published procedure^6^. One microgram of GST-BIK1 or GST-BIK1-Km was incubated with MBP-BSU1, MBP-BSL1, MBP-BSL2, MBP-BSL3, MBP-BSU1-Kelch, MBP-BSU1-PP or MBP-BSU1-S251A in the kinase buffer (20 mM Tris, pH 7.5, 1 mM MgCl2, 100 mM NaCl and 1 mM DTT) containing 100 μM ATP and 10 μCi [^32^P]-γ-ATP at 30 °C for 3 hr. For kinase assay using immunopurified BIK1, total protein was extracted from *BIK1::BIK1-HA* seedlings using immunoprecipitation (IP) buffer (50 mM Tris, pH 7.5, 50 mM NaCl, 0.3 M sucrose, 1% Triton X-100, 1 mM PMSF and 1× protease inhibitor cocktail), centrifuged at 1700g for 5 min. Supernatants were centrifuged at 16,000g for 10 min. The cleared supernatants were incubated with anti-HA antibody (Roche, 3F10) immobilized on Protein A/G agarose beads (Pierce) for 2 hr. The beads were then washed five times with 1 mL of IP washing buffer 1 (50 mM Tris, pH 7.5, 150 mM NaCl, 0.1% Triton X-100) followed by the kinase buffer (25 mM Tris, pH 7.5, 10 mM MgCl2 and 1 mM DTT). Aliquots of BIK1-HA-bead complex were incubated with 1 μg of MBP-BSU1 in the kinase buffer containing 10 μM ATP and 10 μCi [^32^P]-*γ*-ATP at room temperature with shaking at 1,000 rpm for 1.5 hr. Proteins were eluted with 2× SDS sample buffer and separated by SDS-PAGE. Autoradiograph images were obtained by Typhoon Trio imager (GE healthcare).

### Co-immunoprecipitation assays

For transient protein co-expression in *N. benthamiana, Agrobacterium* (GV3101) cells containing *35S::BIK1-Myc* or *35S::BSU1-YFP* expression vectors were washed and resuspended in the induction medium (10 mM MES buffer at pH 5.6, 10 mM MgCl2 and 150 μM acetosyringone) and were infiltrated into young *N. benthamiana* leaves. Two days after infiltration, the infiltrated leaves were harvested. For the MEKK1-BSU1 interaction, *Agrobacterium* (GV3101) cells containing *35S::BSU1-Myc* and *35S::MEKK1-YFP* expression constructs were co-infiltrated in *N. benthamiana* leaves in the same way. Total proteins were extracted with an IP buffer. The proteins were immunoprecipitated using anti-GFP antibody-bead complexes. After washing twice with 0.5 mL of IP washing buffer 2 (50 mM Tris, pH 7.5, 50 mM NaCl, 30 mM Sucrose, 0.2% Triton X-100, 1× protease inhibitor, 0.2 mM PMSF), the co-immunoprecipitated proteins were eluted by 2× SDS sample buffer.

For experiments using *Arabidopsis*, 12-day old wild-type, *35S::BIK1-Myc*, and *35S::BIK1-Myc;BSU1pro::BSU1-YFP* seedlings were crosslinked for 15 min in 1% formaldehyde under vacuum. Total protein was extracted with an IP buffer. After centrifugation at 16,000g for 10 min, the supernatant was incubated with anti-GFP (10 μg) immobilized on UNOsphere SUPrA beads (Biorad) at 4 °C for 2.5 hr. The beads were then washed five times with 1 mL of IP washing buffer 1 and eluted samples were analyzed by immunoblot.

### Quantification of flg22-induced BSU1 phosphorylation

Samples for initial screening for flg22-regulated phosphorylation sites on BSU1 were prepared as the following. Transgenic *35S::BSU1-YFP* seedlings were grown on ^14^N (½ strength MS nutrient without nitrogen (PhytoTechnology Laboratories), NH4NO3 [0.5 g/L, Sigma], KNO3 [0.5 g/L, Sigma], pH 5.7) or ^15^N media (½ strength MS nutrient without nitrogen, ^15^NH ^15^NO [0.5 g/L, Cambridge Isotope Laboratory], K^15^NO3 [0.5 g/L, Cambridge Isotope Laboratory], pH 5.7) for 10 days. ^14^N-and ^15^N-labeled samples were treated with 1 μM flg22 or mock solution for 10 min in reciprocal experiments. Equal amounts of tissues ground in liquid nitrogen were mixed. Proteins were then extracted in an IP buffer with a phosphatase inhibitor (Roche, PhosSTOP), centrifuged at 1700g for 5 min and filtered through miracloth, and centrifuged at 16,000g for 10 min. The supernatant was transferred to a new tube, and the same volume of IP buffer without detergent was added to dilute the detergent to 0.5%, then was incubated with anti-GFP antibody with UNOsphere SUPrA bead (Biorad) at 4 °C for 1 hr, followed by 5 times washing with wash buffer. Samples for targeted quantification of S251 phosphorylation on BSU1 were prepared as the following. The *35S::BSU1-YFP* seedlings were grown on ^14^N or ^15^N media for 13 days and treated with 1 μM flg22 or mock solution for 10 min. Total proteins from mixed samples (^14^N-flg22/^15^N-mock and ^14^N-mock/^15^N-flg22) were extracted with IP buffer 2 with 1× Protease Inhibitor (Pierce) and phosphatase inhibitor (Roche, PhosSTOP), and centrifuged for 1700g for 5 min. Resulting supernatant was centrifuged at 16,000g for 10 min. The supernatant was transferred to a new tube and incubated with anti-GFP nanobody conjugated magnetic agarose beads (Allele Biotechnology) at 4 °C for 1 hr, followed by 5 times washing with IP wash buffer 4 (50 mM Tris, pH 7.5, 150 mM NaCl, 0.05% Triton X-100). Samples were eluted with 2× SDS sample buffer, separated by SDS-PAGE, followed by Colloidal Blue staining (Invitrogen).

BSU1-YFP protein bands were excised and subjected to in-gel digestion with trypsin. Peptide mixtures were desalted using C18 ZipTips (Millipore). Data were acquired in the Data Dependent Acquisition (DDA) mode. Briefly, peptides were analyzed by liquid chromatography– tandem mass spectrometry (LC-MS) on a Nanoacquity ultraperformance liquid chromatography system (Waters) connected to Linear Trap Quadrupole (LTQ) Orbitrap Velos mass spectrometer (Thermo). Peptides were separated using analytical Easy-Spray C18 columns (75 μm × 150 mm) (Thermo, ES800). The flow rate was 300 nL/min, peptides were eluted by a gradient from 2 to 30% solvent B (acetonitrile/0.1%formic acid) over 57 min, followed by a short wash at 50% solvent B. After a precursor scan was measured in the Orbitrap by scanning from mass-to-charge ratio 350 to 1400 at a resolution of 60000, the six most intense multiply charged precursors were selected for collision-induced dissociation (CID) in the linear ion trap.

MS/MS data were converted to peaklist using a script PAVA (peaklist generator that provides centroid MS2 peaklist)^41^, and data were searched using Protein Prospector against the TAIR database *Arabidopsis thaliana* from December 2010 (https://www.arabidopsis.org/), concatenated with sequence randomized versions of each protein (a total of 35386 entries). A precursor mass tolerance was set to 20 ppm and MS/MS2 tolerance was set to 0.6 Da. Carbamidomethylcysteine was searched as a constant modification. Variable modifications include protein N-terminal acetylation, peptide N-terminal Gln conversion to pyroglutamate, and Met oxidation. Subsequent searches were performed to find those peptides modified by phosphorylation. The search parameters were as above, but this time allowing for phosphorylation on serine, threonine, and tyrosine. Assignment of modified peptide was checked manually and is illustrated in Fig. 3. ^15^N-labeled searches were done the same as mentioned above, considering all 20 amino acids are constantly modified by ^15^N labeling. FDR 1% was set for both proteins and peptides. For Quantification, ^15^N labeling efficiency was manually checked. “^15^N labeling” was chosen as a quantitative method using Protein Prospector with automatic adjustment of L:H intensity ratios with labeling efficiency.

For targeted quantification, data was acquired in the Parallel Reaction Monitoring (PRM) mode. Peptides were separated using an EasyLC1200 system (Thermo) connected to high performance quadrupole Orbitrap mass spectrometer Q Exactive HF (Thermo). Peptides were separated using analytical Easy-Spray C18 columns (75 μm × 250 mm) (Thermo, ES802). The flow rate was 300 nL/min, peptides were eluted by a gradient from 3 to 28% solvent B (80% acetonitrile/0.1% formic acid) over 45 min, 28 to 44% solvent B for 15 min, followed by a short wash at 90% solvent B. Samples were analyzed using PRM mode with an isolation window of 1.4 Th. PRM scans were done using 30000 resolutions (AGC target 2e5, 200 ms maximum injection time) triggered by an inclusion list (Supplementary Table 5). Normalized collision energy 27 was used in a higher-energy dissociation mode (HCD). PRM data were analyzed using skyline software^42^.

### Quantitative real-time PCR

Total RNA was extracted from seedlings using the Spectrum Plant Total RNA kit (Sigma). M-MLV reverse transcriptase (Fermentas) was used to synthesize complementary DNA (cDNA) from the RNA. Quantitative real-time PCR (qPCR) was carried out using LightCycler 480 (Roche) and the SYBR Green Master Mix (Bioline). *FRK1* and *At2g17740* primers were previously described^43^. *ROF2* and *CRK13* primers are listed in Supplementary Table 6. *PP2A* was used as an internal reference^44^.

### Data availability

The RNA-seq data that support the findings of this study have been deposited in GEO with the accession code GSE140037. The mass spectrometry proteomics data are available via ProteomeXchange with identifier PXD016283 (Username; Password: pCDR9H2R, for DDA mode) and PXD016257 (Username; Password: IQKH8jha, for PRM mode).

## Acknowledgements

We thank Jian-Min Zhou for sharing *BIK1::BIK1-HA* and *MEKK1-FLAG* seeds, Libo Shan and Ping He for *GST-BIK1* and *GST-BIK1Km* constructs. We thank Andres Valentino Reyes for generating the bee swam box plot figures. This research was supported by grants from NIH (R01GM066258 to Z-Y.W.), National Research Foundation of Korea funded by the Ministry of Science, ICT, Future Planning (NRF-2021R1A2C1006617 and 2020R1A6A1A06046728 to T.W.K., 2021R1A2C1007516 to S-K.K.), NSF (NSF-IOS 2026368 to M.B.M.), NIH

(R01GM135706 to S.L.X.), by Howard Hughes Medical Institute and NIH (P41GM103481 to A.L.B.), and Carnegie endowment fund to the Carnegie mass spectrometry facility.

## Contributions

C.H.P., Y.B., T-W.K. and Z-Y.W. designed the research. C.H.P. and Y.B. performed most of the experiments. C.H.P and Y.B. performed PAMP treatment and immunoblot. C.H.P. performed RNA-seq. C.H.P. and Y.B. analyzed RNA-seq data. C.H.P., Y.B., J-H.Y., S-H.K., S-K.K. and T-W.K. performed and analyzed yeast-two-hybrid assay and *in vitro* pull-down assay. S-H.K., S-K.K. and T-W.K. performed SA quantification. C.H.P., S-L.X, and R.S. performed MS analysis. C.H.P., J-G.K. and M.B.M. performed and analyzed bacterial growth assay. C.H.P., Y.B. and N.Y.X. performed immunoblot and seedling flood inoculation assay. C.H.P., Y.B., S-K.K., T-W.K., M.B.M. and Z-Y.W. wrote the manuscript.

## Additional information

Supplementary Information is available for this paper.

## Supplementary information

Supplementary Table 1. Genes differentially expressed in *bsu-q* vs. wild type (Col-0).

RNA-seq data of genes that are differentially expressed in *bsu*-*q* compared to wild type with fold change >2 (UP: *bsu-q/Col-0*; DOWN: *Col-0/bsu-q*) and *q*-value<0.05.

Supplementary Table 2. GO enrichment analysis of genes differentially expressed in *bsu-q*.

Supplementary Table 3. RNA-seq analysis of flg22 responsive genes in wild type and *bsu-q*. RNA-seq analysis of wild-type (a) and *bsu-q* (b) plants 1 hr after treatment with 1 μM flg22 or mock.

Supplemental Table 4: RNA-seq data of WT and *bsu-q* treated with mock (ct) or 1 μM flg22 (fg) for 1 hr. (a) Data for Fig. 1b. (b) Data for Fig. 1c.

Supplementary Table 5. Inclusion list for S251 PRM.

Supplementary Table 6. Oligos used in this study.

---

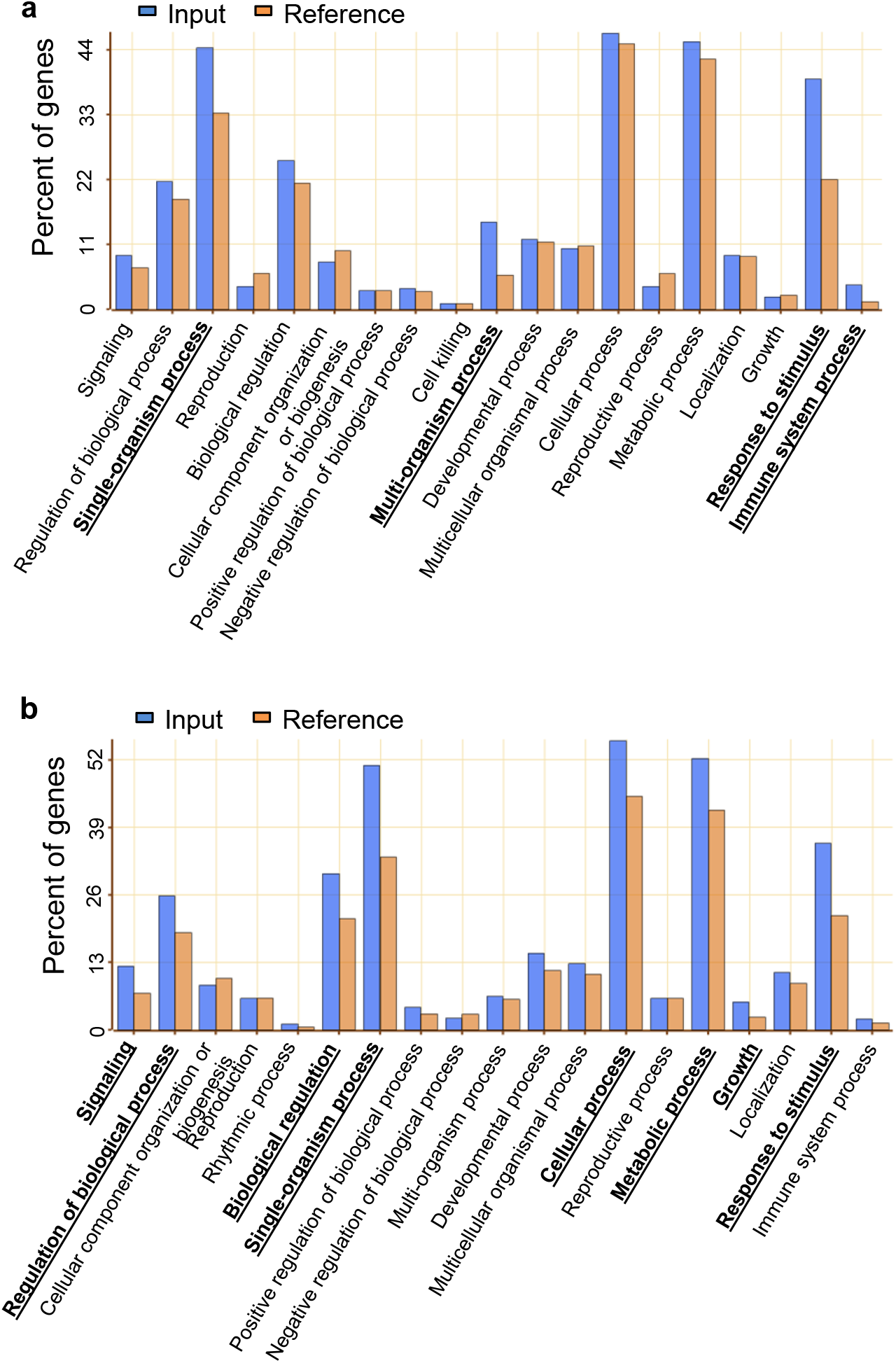

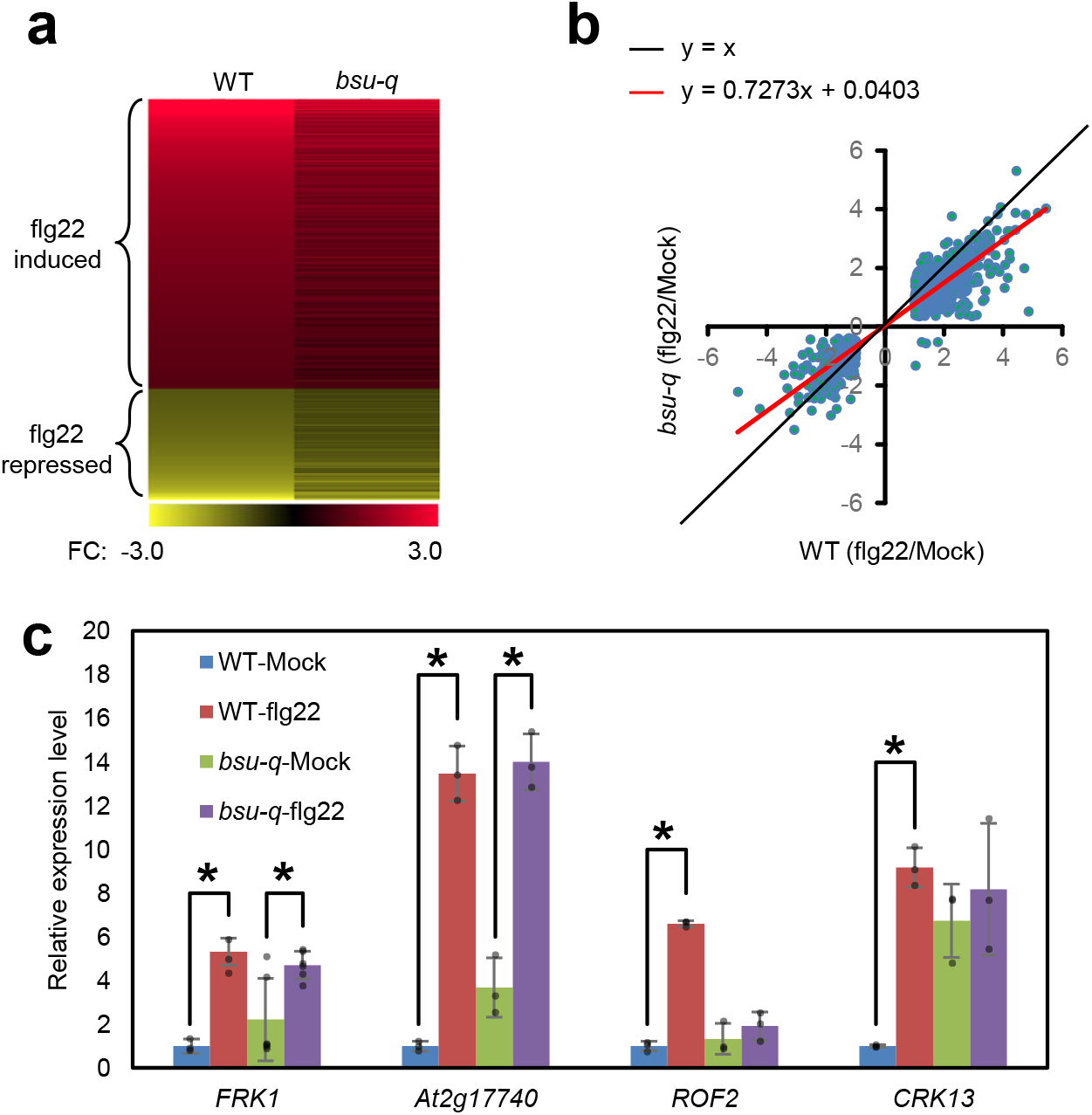

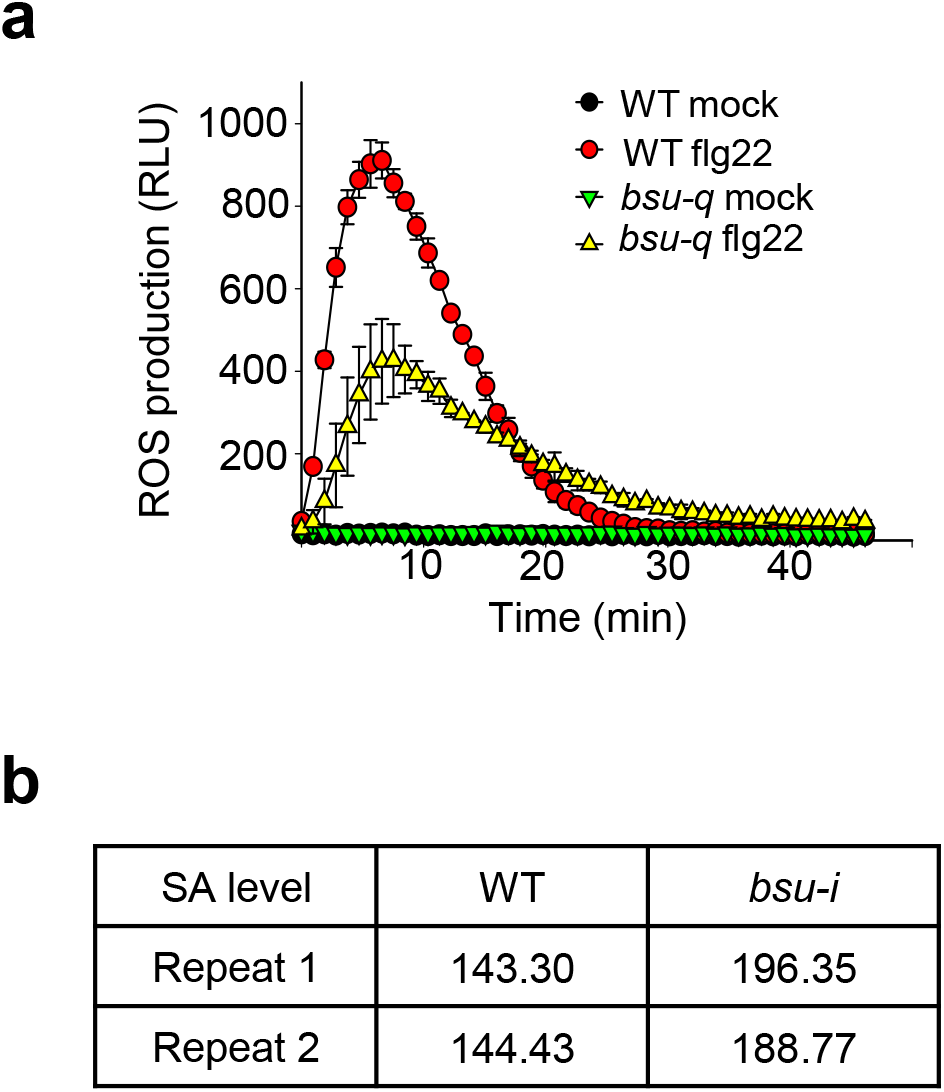

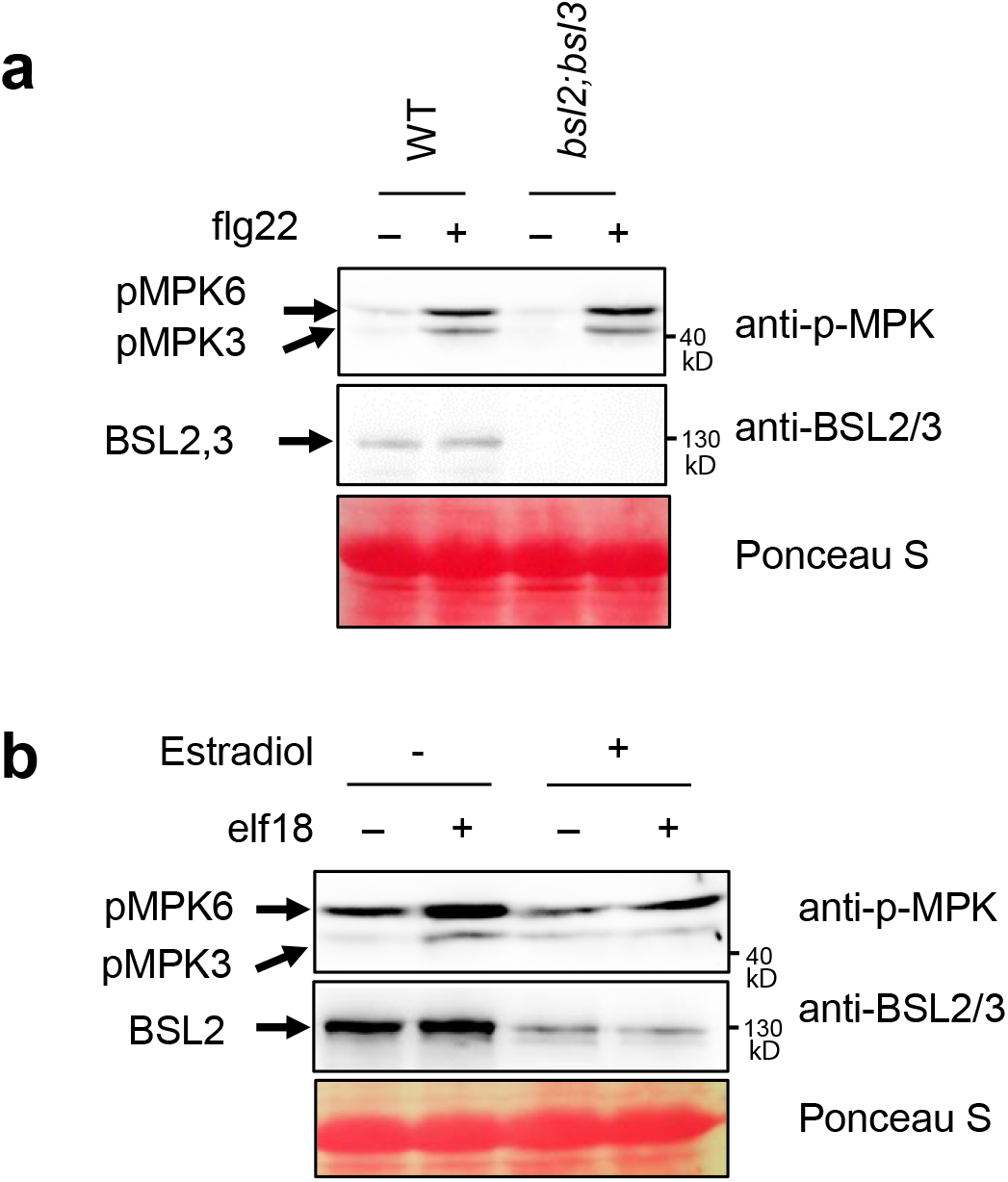

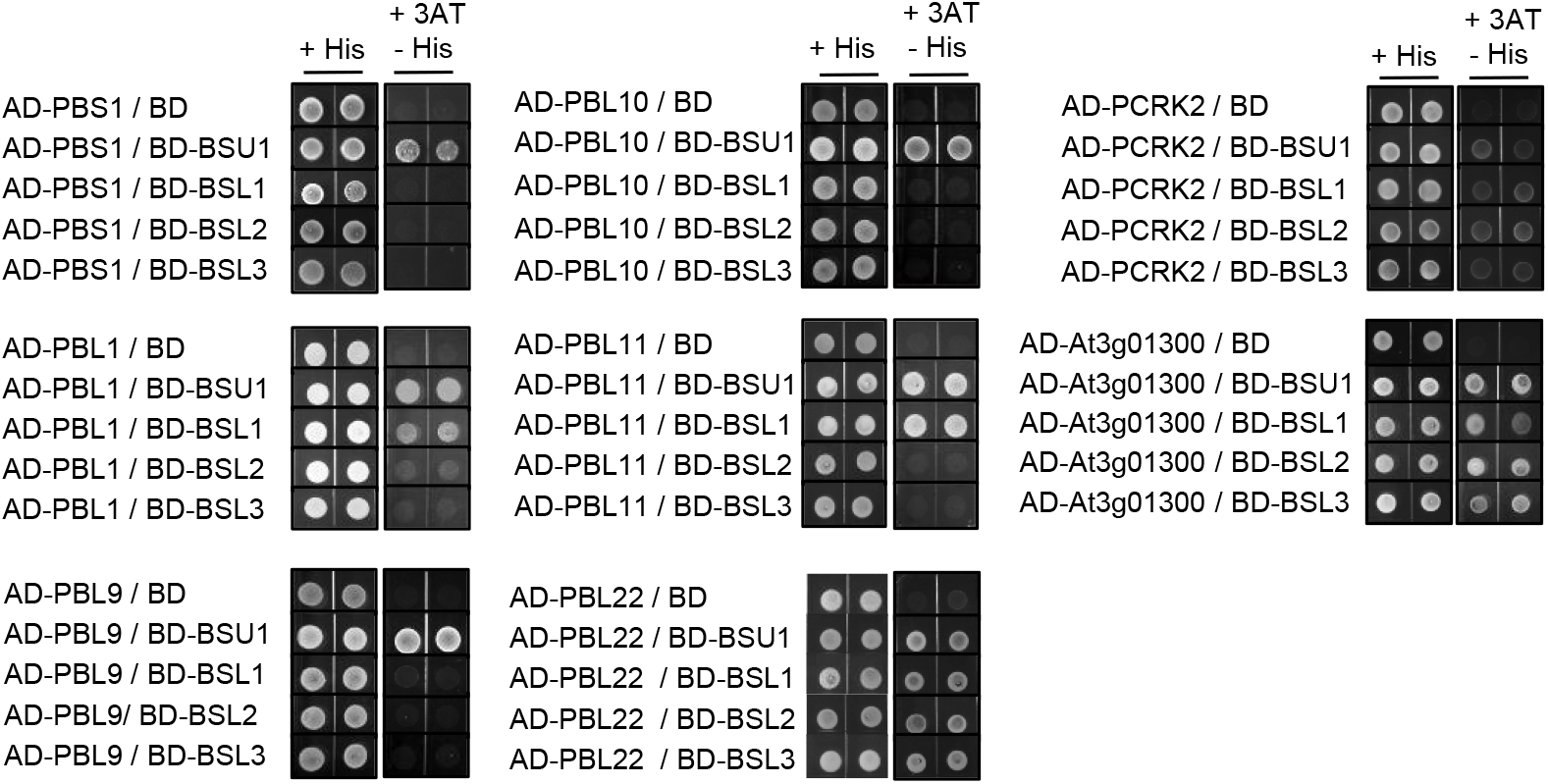

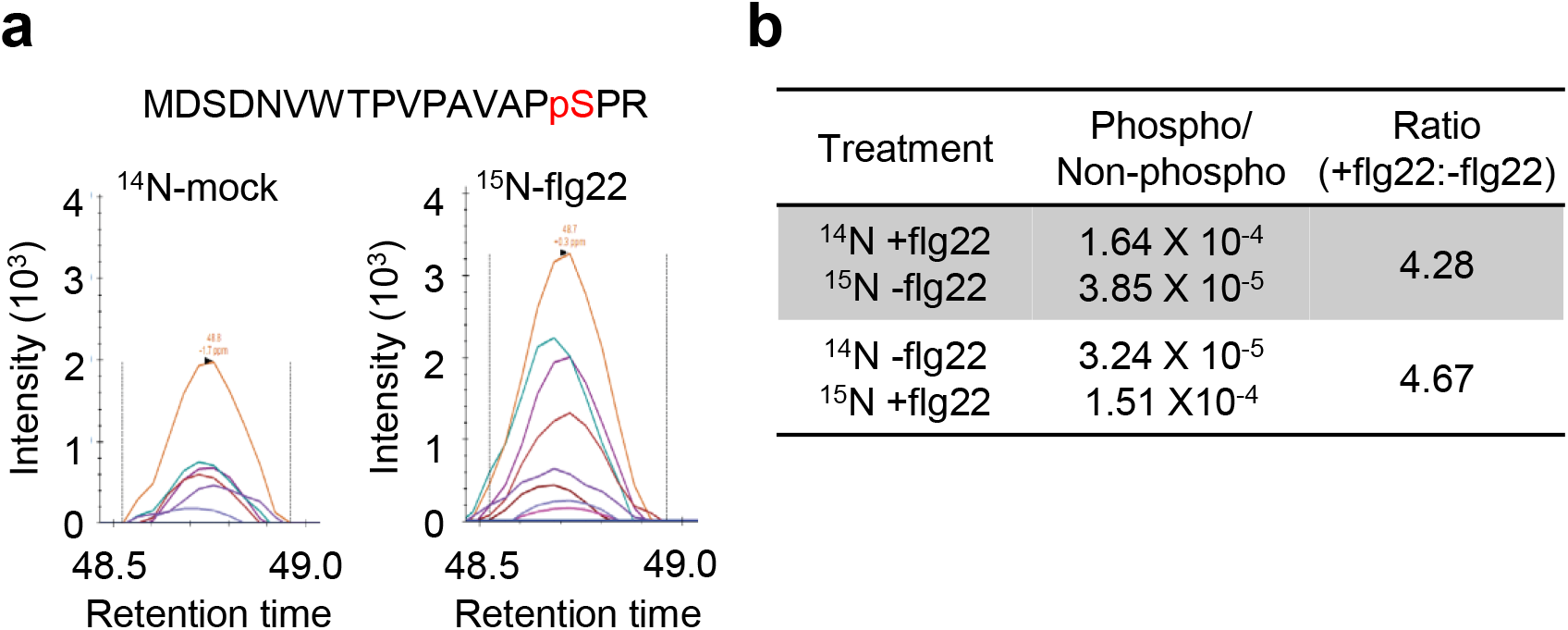

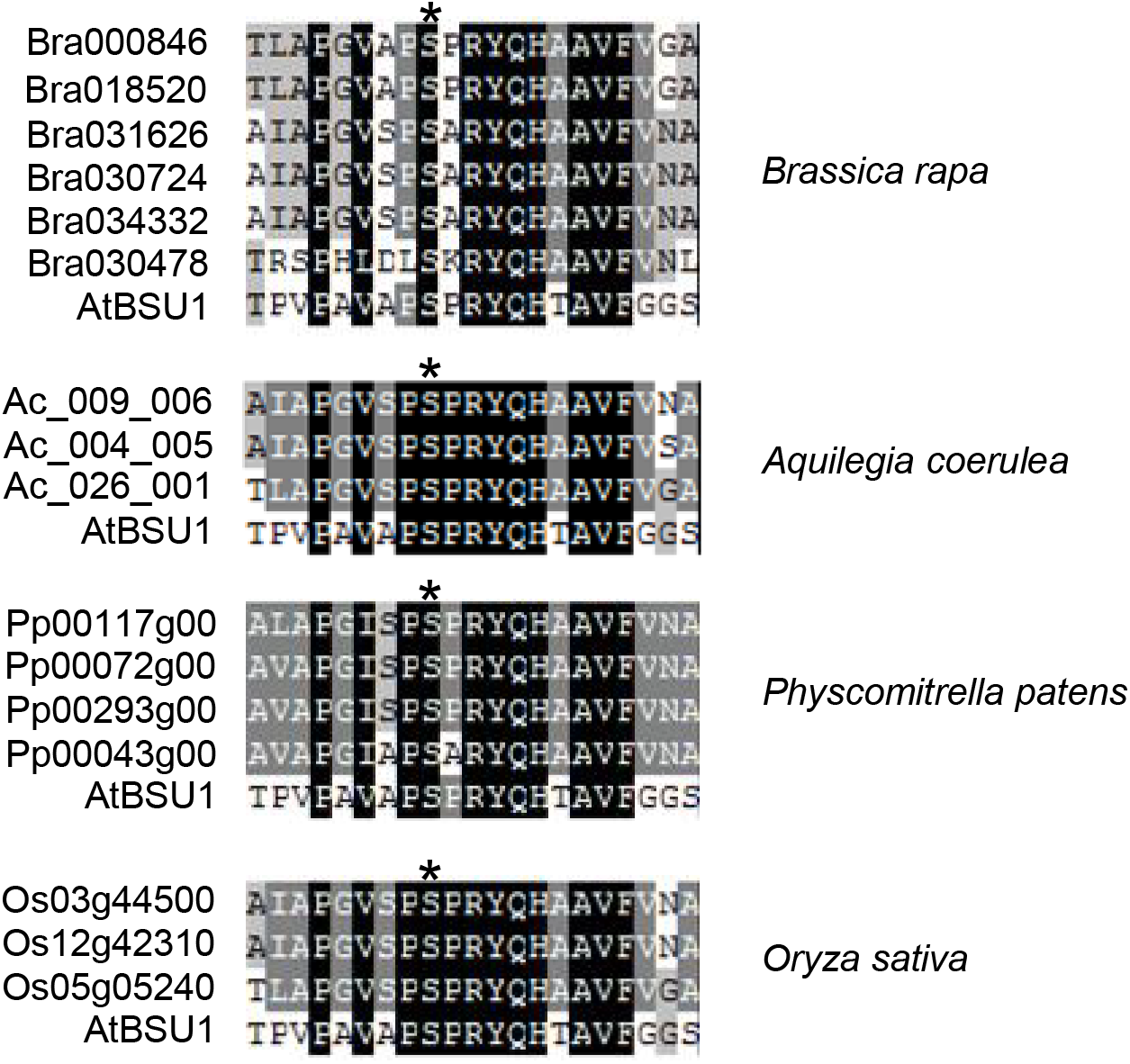

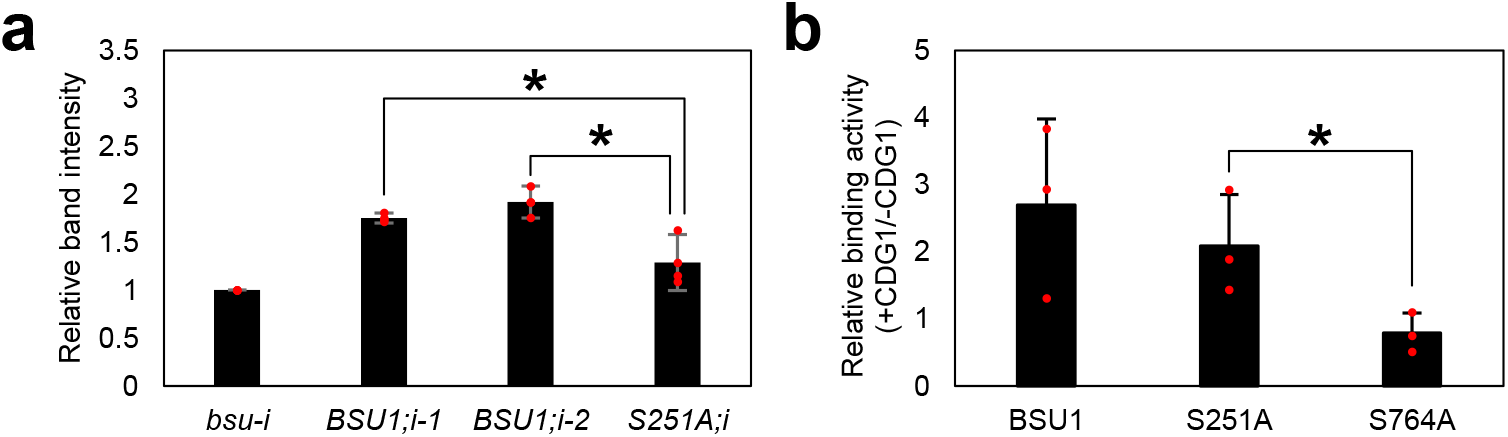

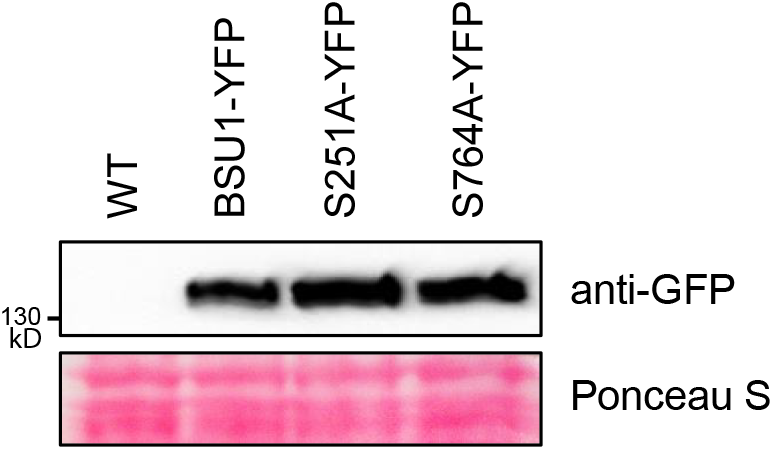

